# Ferritinophagy Contributes to Iron Accumulation and Ferroptosis in Fuchs Endothelial Corneal Dystrophy

**DOI:** 10.64898/2026.08.08.743691

**Authors:** Zachary Shepard, Jessica M. Skeie, Hanna Shevalye, Timothy Eggleston, Linhan Li, Matthew G. Field, Gregory A. Schmidt, Pornpoj Phruttiwanichakun, Christopher S. Sales, Aliasger K. Salem, Mark A. Greiner

**Affiliations:** University of Iowa Carver College of Medicine, Department of Ophthalmology and Visual Sciences, Iowa City, IA 52242; Iowa Lions Eye Bank, Coralville, IA 52241; Minnesota Eye Consultants, Minneapolis, MN 55305; University of Iowa College of Pharmacy, Department of Pharmaceutical Sciences and Experimental Therapeutics, Iowa City, IA 52242

## Abstract

**Purpose:** Fuchs endothelial corneal dystrophy (FECD) is a progressive disease, causing premature death of corneal endothelial cells (CECs). Iron-dependent lipid peroxidation and ferroptosis mediate cell death in FECD. We aimed to determine whether FECD progression is mediated by derangements in ferritinophagy – a form of autophagy that degrades ferritin to release labile ferrous iron – and whether ultraviolet A (UVA) exposure drives FECD progression by activating ferritinophagy.

**Methods:** Endothelium-Descemet membrane (EDM) tissues were collected from patients with end-stage FECD undergoing endothelial keratoplasty and from healthy age-matched donor corneas. Separately, immortalized FECD and healthy control CEC lines were cultured. Cellular levels of NCOA4 production and LC3 activation, both markers of ferritinophagy, were quantified using western blotting and PCR. UVA-exposed immortalized cells were plated on coverslips, stained for immunohistochemistry (IHC), and analyzed using confocal microscopy. Corneal endothelial peels were stained and analyzed using laser ablation–inductively coupled plasma–mass spectrometry (LA-ICP-MS).

**Results:** Surgically explanted FECD CECs showed significantly increased levels of NCOA4 compared to healthy controls. LC3 activation was increased in FECD immortalized CECs; UV exposure further increased LC3 activation. Additionally, UVA exposure showed trends of increased expression of NCOA4 in immortalized FECD and healthy CECs. On IHC of FECD surgical explant tissue, ferritin was decreased markedly, NCOA4 localized in a dramatic punctate pattern, and both ferritin and LC3 localized within cell nuclei. Spectrometry images showed higher iron levels correlating with areas of higher FECD disease burden.

**Conclusions:** Our results demonstrate ferritinophagy in FECD indicated by the increase of NCOA4 and LC3 ferritinophagy markers in FECD patient and cell culture models. Our finding that UVA activates ferritinophagy implicates this mechanism in UVA-mediated FECD progression. Altogether, aberrant iron dysregulation associated with FECD and ferroptosis may be mediated by ferritinophagy, providing a biomarker to assess disease severity as well as a potential target for future medical therapeutics.

## INTRODUCTION

Fuchs endothelial corneal dystrophy (FECD) is a degradative inherited disease with multiple possible causative mutations that result in the permanent death of corneal endothelial cells (CECs) [1]. This disease affects 4% of the population over 40 years of age, making it the most common disorder of the corneal endothelium [2]. As FECD progresses, thickening of Descemet membrane and formation of guttae result in the posterior cornea, which can cause reduced vision due to glare. This, coupled with a decreased density of CECs from cell death related to oxidative damage, can result in progressive corneal edema and vision loss, typically presenting in the fourth or fifth decade [3]. Despite its high prevalence, relatively little is understood about the mechanisms of cell death in the disease process, and corneal transplant remains the only effective treatment because no effective therapeutics have been developed to prevent progression to corneal edema, end-stage disease, and need for surgery [2].

While complete details of FECD molecular pathogenesis are not currently known, aspects of the disease and its progression are increasingly understood. In FECD, the ultimate result of CEC death occurs due to an accumulation of oxidative damage in diseased cells [4]. This damage is reflected in a downregulation of several native antioxidant genes as well as increased amounts of oxidative damage that can severely impact vital organelles including mitochondria and lipid plasma membranes [5]. CECs maintain corneal hydration and transparency through their barrier and pump functions, and as such they have many mitochondria to sustain high levels of metabolic activity [6]. Mitochondrial fusion and fission are necessary to maintain the high volume of mitochondria in CECs, allowing for the selective autophagy of dysfunctional mitochondria in a process called mitophagy. However, this system appears to be dysregulated in FECD, as several studies have demonstrated an increase in mitophagy in CECs in FECD, leading to fewer functioning mitochondria [4, 6, 7]. This has been observed through the utilization of fluorescently tagged LC3, a protein implicated in the process of autophagy [8]. Moreover, other contributors to oxidative damage, such as UV light exposure and specifically UVA which comprises of 95% of solar UV radiation, have been shown to contribute oxidative damage to CECs [5, 9, 10]. Importantly, there is an increasing awareness of the connectivity between FECD genetic background, UVA exposure and cell death due to oxidative damage, and the central role that intracellular iron accumulation plays in mediating cell death resulting from oxidative damage of CEC plasma membranes, a process known as ferroptosis [11].

Ferroptosis is a non-apoptotic form of cell death that results from iron imbalances within the cell [12]. The accumulation of highly reactive iron in its +II oxidative state within the cell leads to the accumulation of intracellular reactive oxygen species (ROS) that are generated by Fenton chemistry reactions. Iron mediated ROS generation damages lipids near the plasma membrane and results in cell death due to plasma membrane rupture. Intracellular iron, which is normally stored in its non-reactive +III oxidative state within the cytosolic protein ferritin, can be released into the cytosol in its reactive +II oxidative state in a process called ferritinophagy [13]. During ferritinophagy, ferritin is tagged by nuclear receptor cofactor 4 (NCOA4) for cytosolic autophagy [14, 15] and Fe^2+^ is released. Ferritinophagy and ferroptosis have been studied and correlated in other diseases, including Parkinson’s disease and autoimmune encephalomyelitis [16–18]; our group has characterized that ferroptosis cell death occurs in FECD [5], but the mechanism of iron accumulation has not been queried. In this study, we investigated the potential role of ferritinophagy in cells with FECD by examining the mRNA transcript, protein, and localization of ferritinophagy markers including ferritin, NCOA4, and LC3-II. Additionally, we compared FECD and non-FECD genotypes as well as examined the results of UV light exposure on these markers to determine whether genetic predisposition and/or UV exposure increases ferritinophagy flux in CECs. While LC3-II serves a role in general cellular autophagy, NCOA4 levels have been specifically implicated in cell-fatal Fe2+ accumulation, with decreased ferritinophagy observed in cells with NCOA4 blockage and knockdown [19]. As such, studying ferritinophagy via NCOA4 levels provided not only a marker of ferritinophagic flux in CECs but also a potential target for the development of future treatments for FECD. Beyond this, we also investigated the association between iron accumulation and FECD activity by examining endothelial cell layers with and without FECD with laser ablation–inductively coupled plasma–mass spectrometry (LA-ICP-MS).

## METHODS

This study was approved by the University of Iowa’s Institutional Review Board (IRB# 201603746) and adhered to the tenants of the Declaration of Helsinki. None of the subjects that provided tissue samples for analysis were from a vulnerable population. All cells and tissues from cornea donors were procured by Iowa Lions Eye Bank; all donors or next of kin provided appropriate consent for tissue donation and research and determined not to constitute human subjects research (IRB# 201802815).

### Cell Culture and Donor Tissues

Immortalized human corneal endothelial cells (HCEC-B4G12) were purchased from Leibniz Institute DSMZ (Braunschweig, Germany), and immortalized FECD human corneal endothelial cells (F35T) were generously provided by Dr. Albert Jun (Johns Hopkins University, Baltimore, MD). F35T cells were originally derived from an FECD patients with a hallmark *TCF4* trinucleotide repeat expansion as described previously [5, 20]. Briefly, both cell lines were cultured in Opti-MEM® I Reduced-Serum Medium (Thermo Fisher Scientific, Waltham, MA, USA) supplemented with 5 ng/mL of human epidermal growth factor (hEGF, Thermo Fisher Scientific), 20 ng/mL of nerve growth factor (NGF, Fisher Scientific), 200 mg/L of calcium chloride (SigmaAldrich, St. Louis, MO, USA), 50 μg/mL of gentamicin (Thermo Fisher Scientific), 50 mg/mL of Normocin^TM^ (Invivogen, San Diego, CA, USA), 0.08% chondroitin sulfate (Sigma-Aldrich), and 8% fetal bovine serum (HyClone Characterized, US origin). Although it is not recommended to grow B4G12 immortalized CECs in high serum per manufacturer recommendations, we chose to maintain identical growth conditions between these two cell lines during the study and opted to use the F35T cell line media conditions. We did not detect morphological transition of B4G12 CECs during the growth period for any of the experiments conducted. Cells were incubated at 37°C with 5% CO_2_ and passaged when confluent.

### Immunohistochemistry

Immortalized CECs were grown on glass coverslips, fixed with 4% paraformaldehyde (PFA), and stored in phosphate-buffered saline (PBS) at 4°C. Human donor corneas and human surgical explants from patients with FECD were collected at the time of surgical resection during endothelial keratoplasty surgery at the University of Iowa and fixed in 4% PFA for 10 minutes prior to storage at 4°C in PBS – donor and surgical explant tissues were treated as whole peels during IHC experimentation. Patient demographics are listed in **Table 1**. Cells were labeled with primary antibodies targeted against LC3-II (catalog# 18722-1-AP, lot# 00124153, Proteintech, Rosemont, IL, USA), NCOA4 (catalog# AM2019a, lot# CA111220D, ABCEPTA, San Diego, CA, USA), and ferritin (catalog# PA1-29381, lot# ZD4287375E, Invitrogen, Thermo Fisher Scientific) for colocalization assays. Secondary antibodies were targeted to each antibody species: LC3-II (Alexa Fluor 488, Anti-Rabbit IgG, catalog# 111-547-003, lot# 159615, Jackson ImmunoResearch, West Grove, PA, USA); NCOA4 (Alexa Fluor 660, Anti-Mouse IgG, catalog# A21055, lot# 2720370, Invitrogen); and ferritin (Alexa Fluor 594, Anti-Rabbit IgG, catalog# 111-587-003, lot# 161439, Jackson ImmunoResearch). A specific fluorescence channel (red, green, or indigo) was dedicated to ferritin, LC3-II, and NCOA4, respectively; these channels were merged to observe their relative positions in the CECs as an indicator of ferritinophagic flux. For nuclear counterstain, samples were incubated with Mayer’s hematoxylin (catalog# 26043-05, Electron Microscopy Sciences, Morgantown, PA, USA) for 4 minutes followed by 1 minute rinse with alkaline distilled water (pH 9.0). Then samples were rinsed with abundance of non-alkaline distilled water, flat mounted on glass slide and air dried at 37C for 48 hours. Whole sample bright field scans were collected by Olympus BX63F microscope (Olympus Surgical Technologies America, Bartlett, TN, USA) under control of multi-dimensional image acquisition algorithm in cellSens software (version 3.2, Evident Corporation, Nagano, Japan).

**Table 1.**
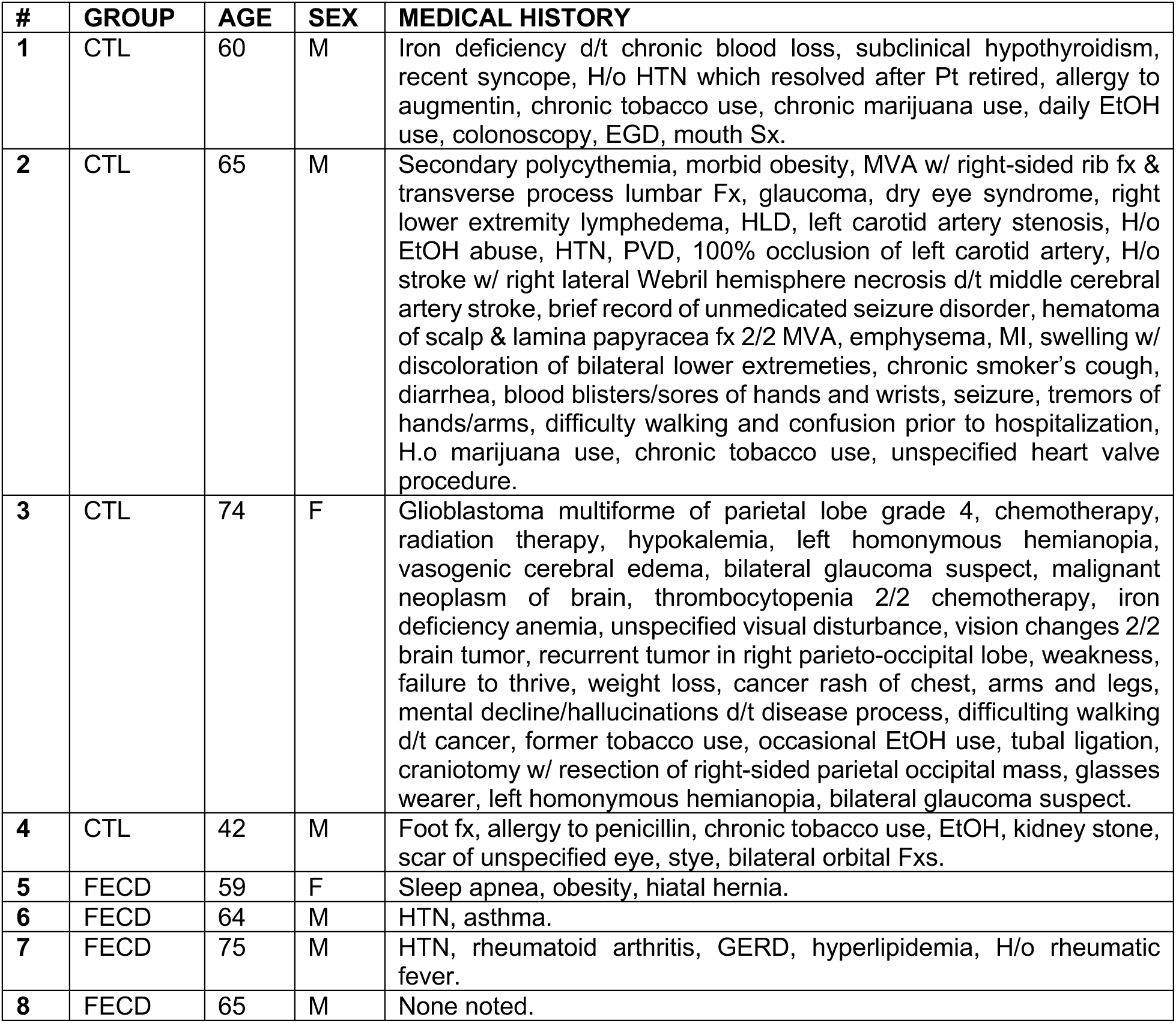
Demographic and medical history information for keratoplasty patients with FECD and human donor tissues without surgical history (controls; CTL) used for immunohistochemistry. Abbreviations: AAA, abdominal aortic aneurysm; AD, advanced diabetes; A-fib, atrial fibrillation; CABG, coronary artery bypass grafting; CAD; coronary artery disease; CHF, congestive heart failure; CKD, chronic kidney disease; COPD, chronic obstructive pulmonary disease; DJD, degenerative joint disease; DVT, deep vein thrombosis; ESRD, end stage renal disease; GERD, gastroesophageal reflex disease; HLD, hyperlipidemia; HTN, hypertension; hx, history; IDDM, insulin dependent type II diabetes mellitus; MI, myocardial infarction; NAD, nonadvanced diabetes; NASH, non-alcoholic steatohepatitis; ND, nondiabetic control; NIDDM, non-insulin dependent type II diabetes mellitus; OSA, obstructive sleep apnea; PONV, post-operative nausea and vomiting; PSA, prostate-specific antigen; PVD, peripheral vascular disease; RA, rheumatoid arthritis; SOB, shortness of breath; STEMI, ST-segment elevation myocardial infarction; sx, symptoms; TURP, transurethral resection of the prostate; UTI, urinary tract infection.

### Quantitative PCR

mRNA was collected from cell and donor tissue samples using TriZOL (ThermoFisher). Transcripts were reverse transcribed; cDNA was amplified using standard PCR cycling protocols with primers designed for *NCOA4, FTH1, FTMT,* and *FTL* and normalized to the housekeeping gene, *18S*. Quantities were compared between FECD and control groups using Student’s t-test. Primers used are listed in **Table 2**. For PCR experiments, fold change data is reported in text, while graphs include RQmin and RQmax error bars.

**Table 2.**
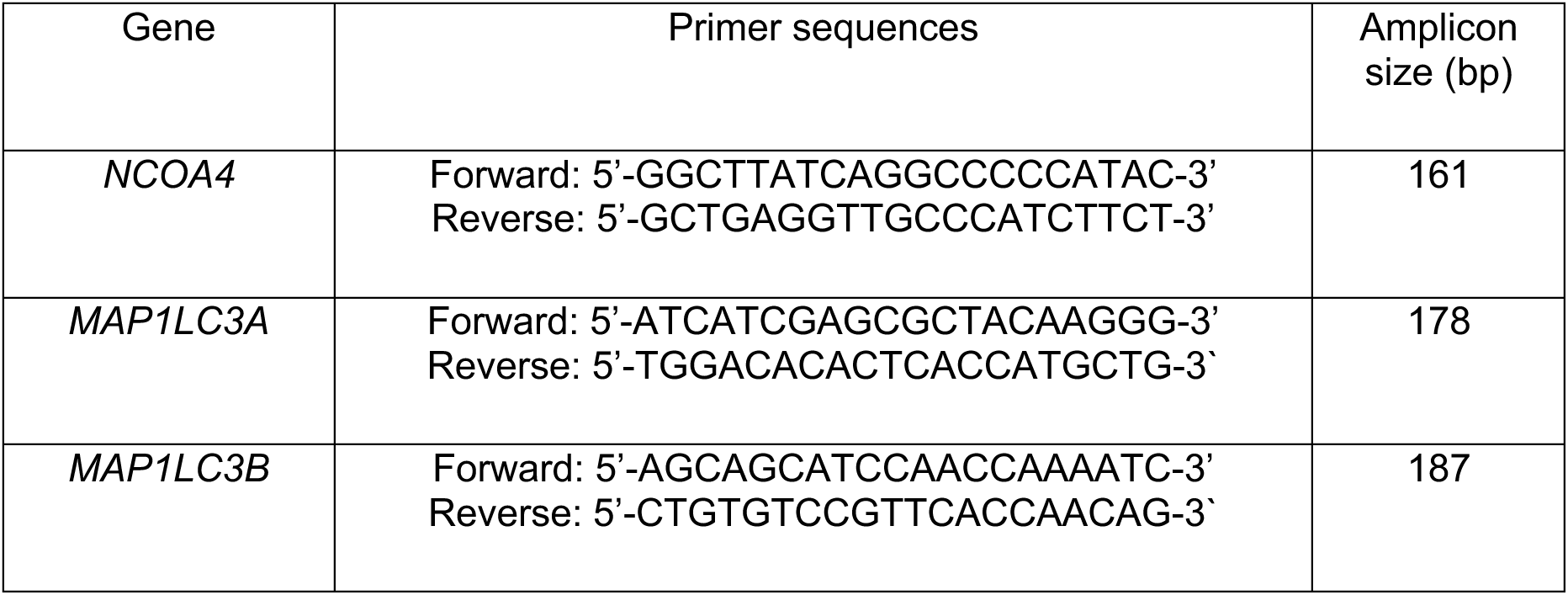
List of primers used for PCR experiments.

### Automated Microfluidic Western Blotting

Proteins from cells and human corneal tissues were lysed using radioimmunoprecipitation assay (RIPA) buffer and quantified using BCA assay. Equal amounts of protein were loaded onto the capillaries of the microfluidic western blotting cartridge for the Jess Western instrument (Protein Simple, San Jose, CA, USA). Samples were probed for NCOA4, LC3-II, LC3, and ferritin, as well as total protein. Protein quantities were normalized to total protein and compared between FECD and control groups. Statistical analyses were performed using Student’s t-test. Traditional western blot technique was used in the analysis of LC3-II protein levels.

### Oxidative Stress Induction

HCEC-B4G12 and F35T were cultured until confluent in 6-well plates. Media was removed from wells and was replaced with media with various concentrations of antimycin A (AMA). Cell cultures were incubated in AMA-infused media for two hours, after which they were harvested for mRNA and protein. Concentrations ranging from 20uMol to 100uMol AMA were initially used based off cell death rates noted in previous publications [21–23]; this gradient was expanded to include 200, 300, and 500 uMol AMA when we observed insufficient oxidative damage at lower AMA concentrations in CEC cultures.

### UVA Irradiation

HCEC-B4G12 and F35T were cultured in a 24 well plate, 400,000 cells per well. Cells were exposed to UVA irradiation at the fluence of 5 J/cm^2^ and 10 J/cm^2^ using Rayonet Photochemical Reactor (RPR-200, The Southern NE Ultraviolet Co., Brandford, CT, USA). Briefly, four RPR-3500A UVA broadband lamps of 12-inch length emitting 350 nm light with an irradiance of 7.13 mW/cm^2^ were placed at 10 cm above the cells and illuminated 11 min 41 sec to achieve 5 J/cm^2^ UVA, or were illuminated 23 min 22 sec to achieve 10 J/cm^2^ UVA.

#### siRNA-mediated knockdown of NCOA4 in HCEC-B4G12 and F35T cells

A set of three distinct 27-mer siRNA duplexes against human *NCOA4* (Cat#SR305344; referred to as siRNA-A, siRNA-B, and siRNA-C) as well as a universal 27-mer scrambled siRNA duplex (Cat#SR30004; referred to as (-)siRNA) were purchased from Origene Technologies (Rockville, MD, USA). Lipofectamine™ RNAiMax (Cat#13778150; ThermoFisher) was used as the transfection agent for the reverse transfection of HCEC-B4G12 and F35T cells with the anti-*NCOA4* siRNAs and the (-)siRNA control according to the manufacturer’s protocol. The anti-*NCOA4* siRNAs and the (-)siRNA control were reverse-transfected into HCEC-B4G12 (16,500 cells/well/0.2 mL) or F35T (8,250 cells/well/0.2 mL) cells in 96-well plates at a final siRNA concentration of 10 nM. The culture media were renewed at 48 h and the cell lysates were harvested at 72 h after the reverse transfection. To harvest the cell lysates, the cells were lysed using 1X RIPA lysis buffer (Cat#R0278; Sigma-Aldrich, St. Louis, MO, USA) supplemented with a protease inhibitor cocktail (Cat#11836153001; Roche Diagnostics, Mannheim, Germany). NCOA4 protein expression levels in the cell lysates were analyzed using a Jess™ Simple Western™ automated western blot system (bio-techne, Minneapolis, MN, USA) using mouse anti-human NCOA4 polyclonal antibodies (Cat#AM2019a; Abcepta, Inc., San Diego, CA, USA) and anti-mouse secondary HRP antibodies (Cat#042-205; bio-techne) for chemiluminescence detection. The anti-NCOA4 antibody dilutions were 1:20 and 1:80 for the probing of HCEC-B4G12 and F35T cell lysates, respectively. The target NCOA4 signals were normalized to the total protein in each sample using chemiluminescence-based total protein detection kits (Cat#DM-TP01; bio-techne) and RePlex™ modules (Cat#RP-001; bio-techne). The automated western blot data were processed on ProteinSimple Compass software (bio-techne). Based on our findings, the anti-NCOA4 siRNA-C and siRNA-B were selected for NCOA4 knockdown in subsequent experiments for HCEC-B4G12 and for F35T cells, respectively.

#### RSL3-mediated ferroptosis assay in NCOA4-knockdown HCEC-B4G12 and F35T cells

HCEC-B4G12 and F35T cells were reverse-transfected with anti-NCOA4 siRNA-C (for HCEC-B4G12), siRNA-B (for F35T), or the (-)siRNA control using previously mentioned procedures at a final siRNA concentration of 10 nM. At 48 h after transfection, cells were treated with either 5-µM ferrostatin-1 (Fer-1, Cat#SML0583; Sigma-Aldrich) or DMSO (equivalent to the DMSO content in the Fer-1 treated group). The Fer-1-treated groups serve as the anti-ferroptosis control. Fer-1 or DMSO-treated cells were cultured for another 24 h (equivalent to 72 h post-transfection) before being challenged with a ferroptosis-inducing agent, RSL3 (Cat#SML2234; Sigma-Aldrich) at the final concentrations of 0.1, 0.5, or 1.0 µM in complete culture media. Non-RSL3-treated control groups were included and instead treated with DMSO (equivalent to the DMSO content in the 1.0 µM RSL3 group). At 24 h after the RSL3 challenge, cell viability was quantified using an MTS assay (Cat#G3580; Promega, Madison, WI, USA) per manufacturer’s instructions. The MTS assay signals were measured using a SpectraMax™ M5 microplate reader (Molecular Devices, Sunnyvale, CA), and the relative cell viability data were obtained by comparison to the DMSO-treated controls.

#### UVA-induced intracellular ferrous ion (Fe^2+^) changes in NCOA4-knockdown F35T cells

To explore the relationship between the NCOA4 levels and the UVA-induced Fe^2+^ changes, F35T cells (50,000 cells/well/0.5 mL in 24-well plates) were reverse-transfected with the anti-NCOA4 siRNA-B using Lipofectamine™ RNAiMax at a final siRNA concentration of 10 nM. At 72 h after reverse transfection, the cell monolayers were washed twice with Hank’s Balanced Salt Solution (HBSS, Cat#14025076; ThermoFisher) and maintained in HBSS (0.5 mL/well) before UVA exposure. Similar to our prior work, the cell monolayers were exposed to UVA irradiation at a fluence of 5 J/cm^2^ over 700 seconds using a Rayonet Photochemical Reactor (RPR-200, The Southern NE Ultraviolet Co., Brandford, CT) [5]. The UVA-exposed cells were harvested via trypsinization, washed twice using the Live Cell Image Solution (Cat#A59688DJ; ThermoFisher), and stained for intracellular Fe^2+^ using FerroOrange fluorescent probe (Cat#F374; Dojindo Laboratories, Inc., Kumamoto, Japan) per the manufacturer’s protocol at a final concentration of 1 µM in the Live Cell Image Solution. The FerroOrange-stained cells were then analyzed using a flow cytometer (BD FACSCalibur™; BD Biosciences, Franklin Lakes, NJ, USA). The flow cytometry data were analyzed using FlowJo software (BD Biosciences).

### Ferritinophagy activity

Cells and respective controls were analyzed for ferritinophagy markers and flux using qPCR, western blotting, and immunohistochemistry. Because NCOA4 levels have been specifically implicated in ferritinophagic activity [19], heightened protein expression of NCOA4 was utilized as a marker of recent or current ferritinophagy in CECs. Protein levels of LC3 were used to evaluate autophagic activity in CECs. During formation of the autophagosome in ferritinophagy, LC3 is cleaved to form LC3-I; upon autophagy initiation, LC3-I binds to the phagophore membrane to form LC3-II [24]. Because LC3-II remains bound to the autophagosome, it serves as a marker of active autophagy. To assess the relative amount of ongoing autophagy in CECs, the quantity of LC3-II was compared with total LC3 expressed by CECs. This ratio of activated LC3-II over total LC3 was utilized to assess differences in autophagic (and by extension, ferritinophagic) flux between CECs. Previous studies examining ferritinophagy via IHC have fluorescently tagged some combination of NCOA4, LC3, and ferritin, revealing a variety of potential observable patterns of ferritinophagy. Punctate colocalization of LC3 and ferritin has demonstrated ferritinophagy in mouse pulmonary epithelial cells after sepsis-induced lung injury [17]. Alternatively, a different pattern demonstrating ferritinophagy has been observed in mice models of experimental autoimmune encephalomyelitis; IHC images showed tissues in which cells with high NCOA4 fluorescence expressed little ferritin fluorescence (demonstrating recent ferritinophagy), while cells with low NCOA4 expression had high ferritin fluorescence, implying absence of ferritinophagy [18]. We examined CECs stained in triplicate for NCOA4, LC3, and ferritin. Colocalization of these markers in a punctate pattern was interpreted as active ferritinophagy, while high NCOA4 expression with decreased ferritin expression was viewed as a result of recent ferritinophagic activity.

### Iron elemental Imaging by LA-ICP-MS

Following tissue preparation (described above), the spatial distribution of iron across the corneal endothelial layer was mapped by laser ablation–inductively coupled plasma–mass spectrometry (LA-ICP-MS). Ablation was performed with an Elemental Scientific Lasers imageGEO193 laser-ablation system (193 nm) coupled to an Agilent 7800 quadrupole ICP-MS. Tissue sections were ablated line-by-line with a 15 µm laser spot at a repetition rate of 75 Hz, a scan speed of 225 µm s⁻¹, and a fluence of 5.74 J cm⁻²; consecutive line scans were spaced at 15 µm to match the spot size, and within each line the spot advanced 3 µm per pulse (225 µm s⁻¹ at 75 Hz), corresponding to 5-fold oversampling, with a nominal lateral resolution of 15 µm. The ablated aerosol was carried to the ICP in helium, delivered as a cell/chamber flow of 250 mL min⁻¹ and a cup flow of 300 mL min⁻¹. ⁵⁶Fe and ¹³C were acquired without a collision/reaction gas (no-gas mode) to maintain the high acquisition speed required for imaging; the dry ablation aerosol minimizes the ⁴⁰Ar¹⁶O⁺ polyatomic interference on ⁵⁶Fe. To account for point-to-point variation in ablated mass and section thickness, the ⁵⁶Fe signal was normalized to ¹³C and expressed as the ⁵⁶Fe/¹³C intensity ratio, with ¹³C serving as an internal standard on the assumption of a uniform carbon distribution across the section [25]. Iron distributions are therefore reported as semi-quantitative, ¹³C-normalized signal-intensity maps rather than absolute concentrations. Elemental images were reconstructed and normalized in Iolite [26, 27].

### Statistical Analysis

All data were expressed as the mean ± standard error of the mean (SEM) or mean ± standard deviation (SD). Statistical analysis was performed using the two-tailed Student’s t-test when the experimental group was only compared with the control group. One-way ANOVA followed by Tukey’s post-hoc test was utilized when multiple groups were compared with each other. P-values of less than 0.05 were considered statistically significant. All experiments were carried out with at least 3 biological replicates and in technical duplicate. For the western blotting of the post-knockdown NCOA4 protein expression, the mean fold changes of NCOA4 protein expression levels (normalized from total protein-corrected NCOA4 band areas to those of the (-)siRNA control group) were compared between each anti-NCOA4 siRNA sequence (siRNA-A, -B, or -C), and the (-)siRNA controls using ordinary one-way ANOVA with Dunnett’s multiple comparison with the (-)siRNA control (significance level = 0.05). For the RSL3-mediated ferroptosis assays in NCOA4-knockdown cells, the relative cell viability data were compared within each RSL3-treated group of the same RSL3 concentration. In each comparison, the mean %cell viability of the (-)siRNA group was compared to that of the anti-NCOA4 siRNA group using two-tailed unpaired Student’s t-test (significance level = 0.05). For the UVA-induced intracellular ferrous ion changes in NCOA4-knockdown cells, the relative intracellular ferrous ion levels (normalized from median fluorescence intensities (FL2-H) to those of the (-)siRNA control group) were compared between the (-)siRNA control and the anti-NCOA4 siRNA group within the same UVA exposure status (no UVA or 5 J/cm^2^ UVA) using two-tailed unpaired Student’s t-test (significance level = 0.05).

## RESULTS

### *TCF4* FECD genetic background increases production of ferritinophagy markers

To compare ferritinophagic flux between healthy CECs and FECD CECs, protein and mRNA levels of NCOA4 and LC3 were measured as indicators of ferritinophagy in immortalized healthy CECs, transformed immortalized FECD CECs, healthy donor CECs, and surgical explant cells with FECD. Surgical explant FECD cells had 2.2-fold higher levels of NCOA4 protein compared to healthy cornea cells (p<0.01) (**Fig. 1A**). The same pattern was seen in immortalized FECD cells, which showed an 42.1% increase in NCOA4 protein levels compared to healthy immortalized corneal cells (p<0.05) (**Fig 1B**). Additionally, immortalized FECD cells displayed increased ratio of activated LC3B-II protein to total LC3 protein of 0.52 when compared with the LC3 activation ratio of 0.35 in healthy immortalized control cells (p<0.01) (**Fig 1C**). When examining cells via qPCR, surgical explant FECD cells showed decreased NCOA4 mRNA levels – mRNA production was 0.43-fold compared to healthy donor corneal cells (p<0.01) (**Fig 1D**). Immortalized FECD cells showed similar levels of NCOA4 mRNA levels when compared to healthy immortalized corneal cells (**Figure 1E)**. Similarly, immortalized FECD cells similarly showed comparable levels of LC3B mRNA compared with healthy immortalized cells, with 1.07-fold production of LC3B mRNA (**Fig 1F)**. LC3A mRNA levels greatly increased in immortalized FECD cells, with a 379-fold production of LC3A mRNA compared with immortalized healthy cells (**Fig 1G**).

**Figure 1.**
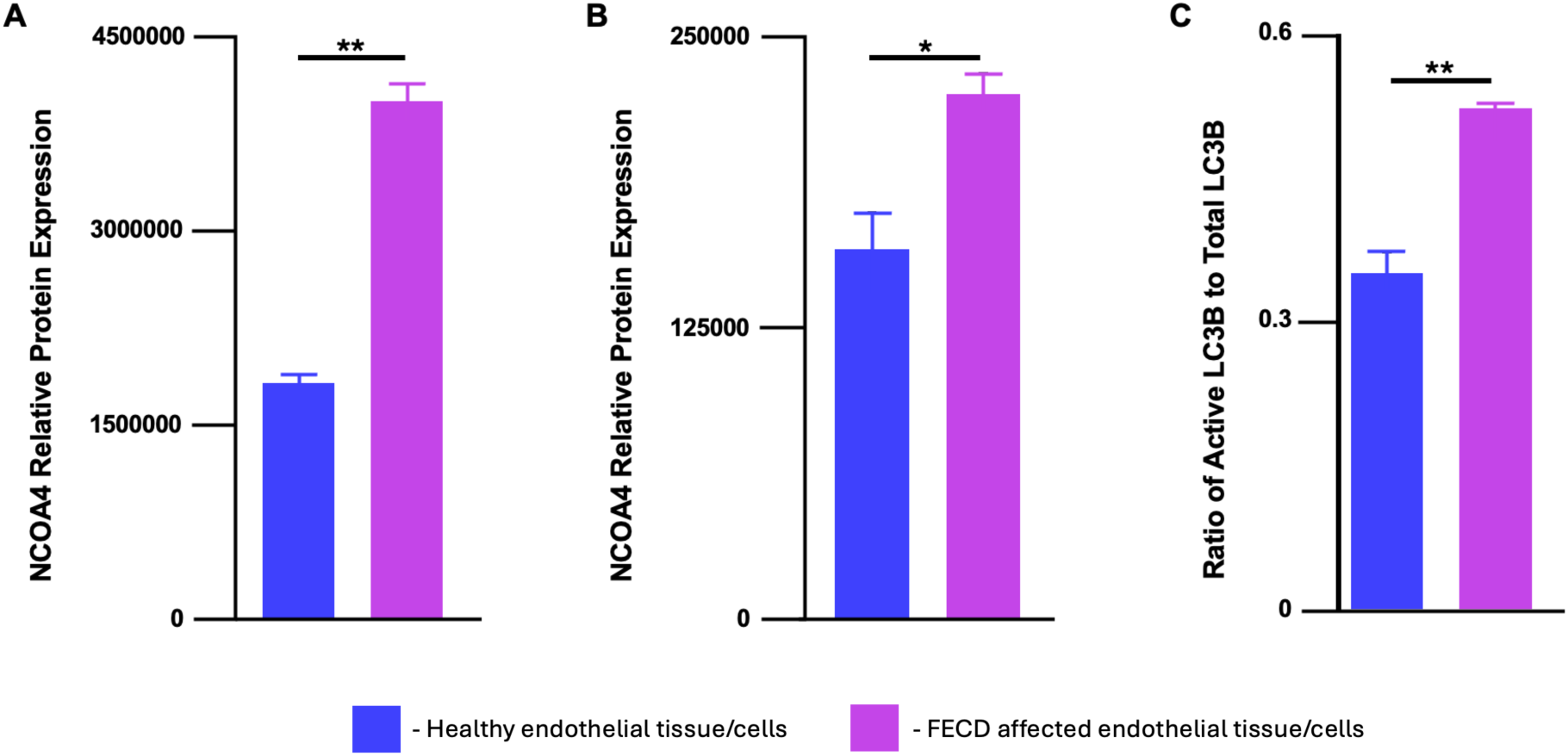
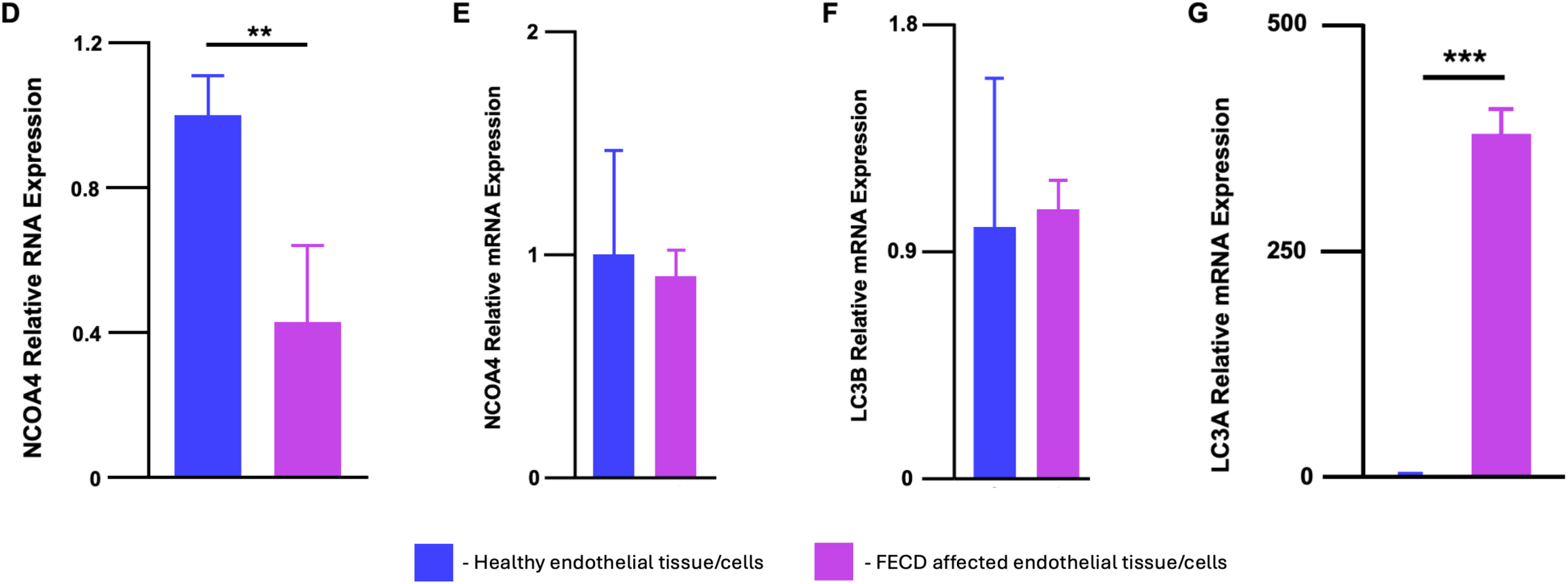
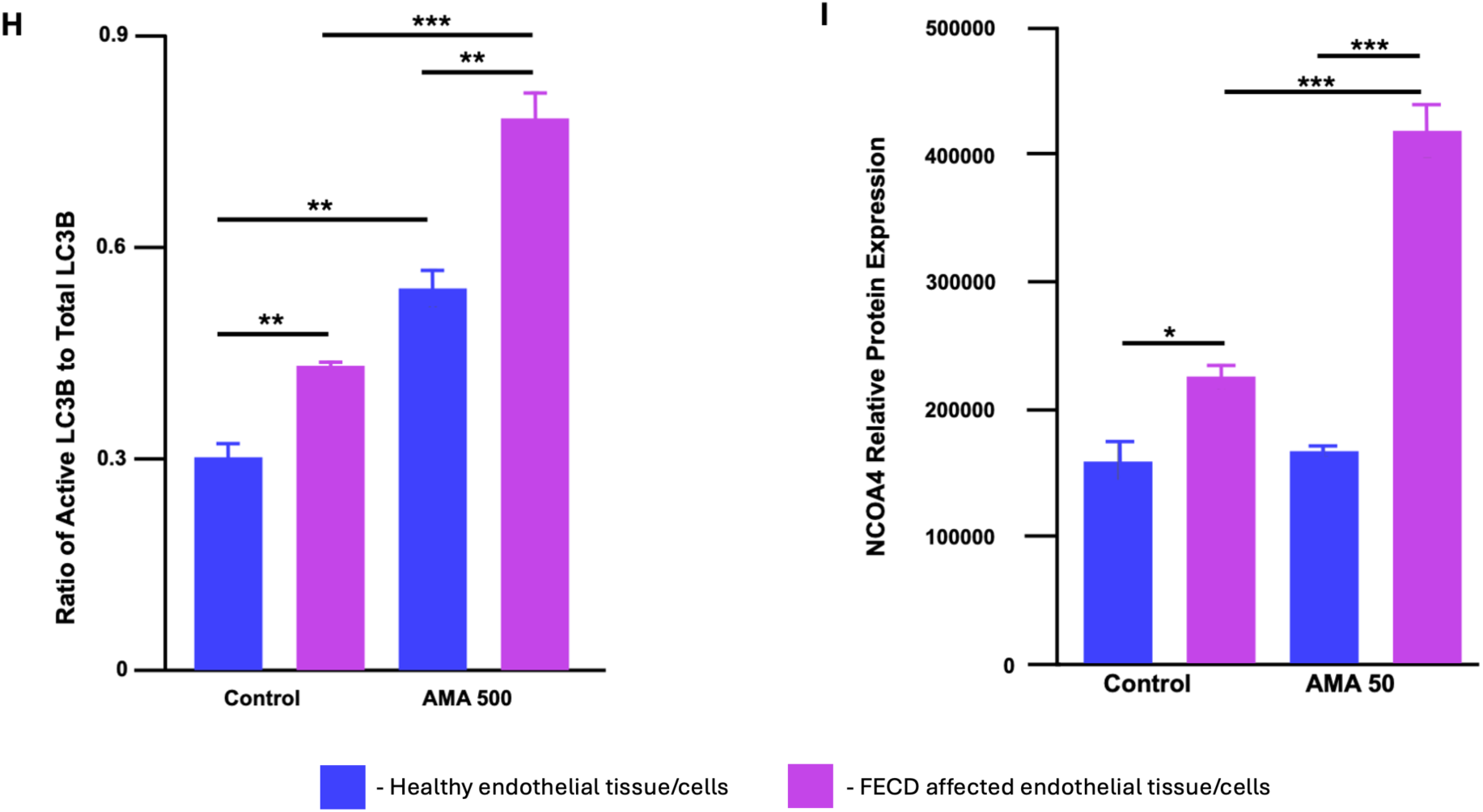
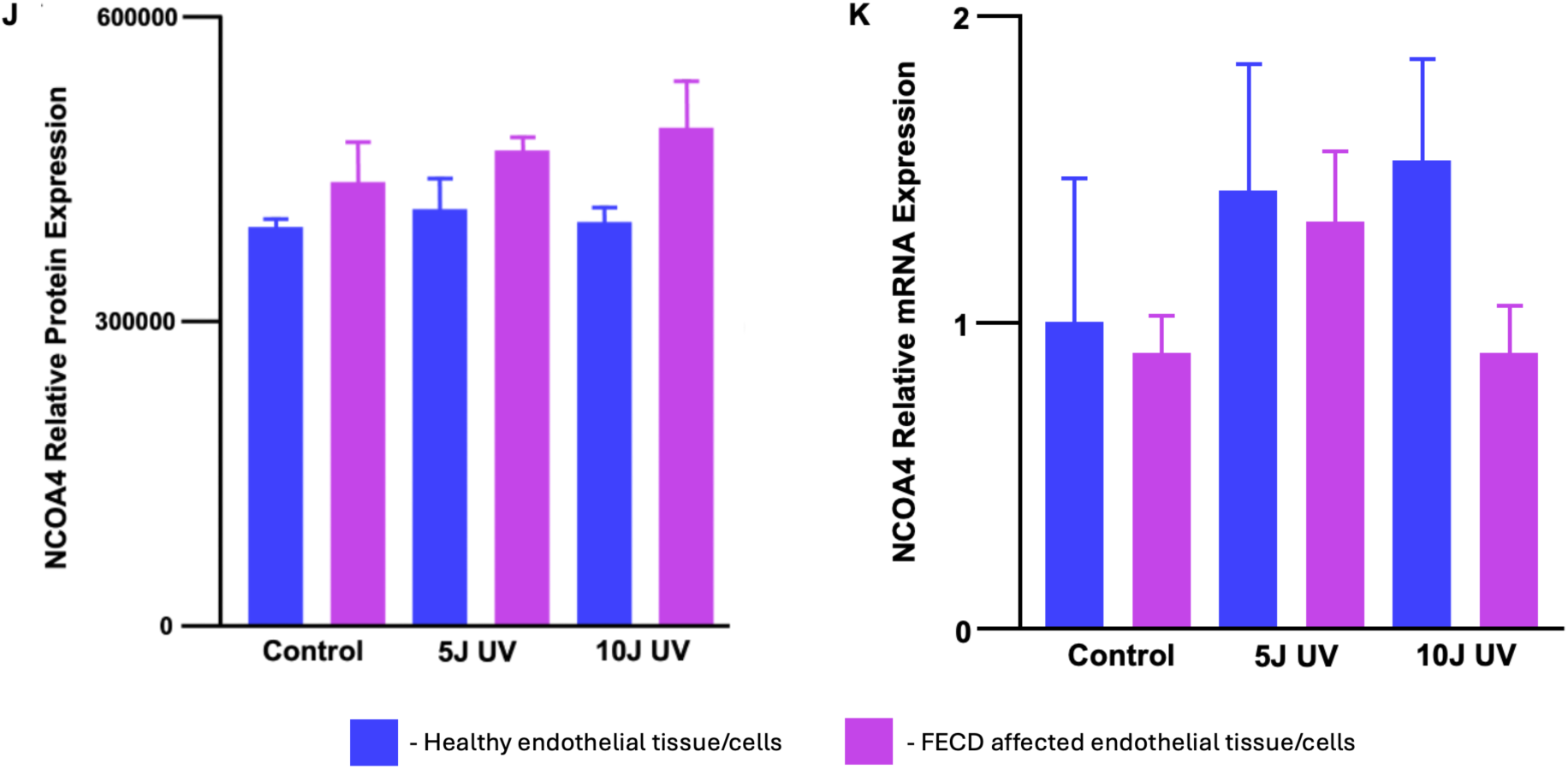
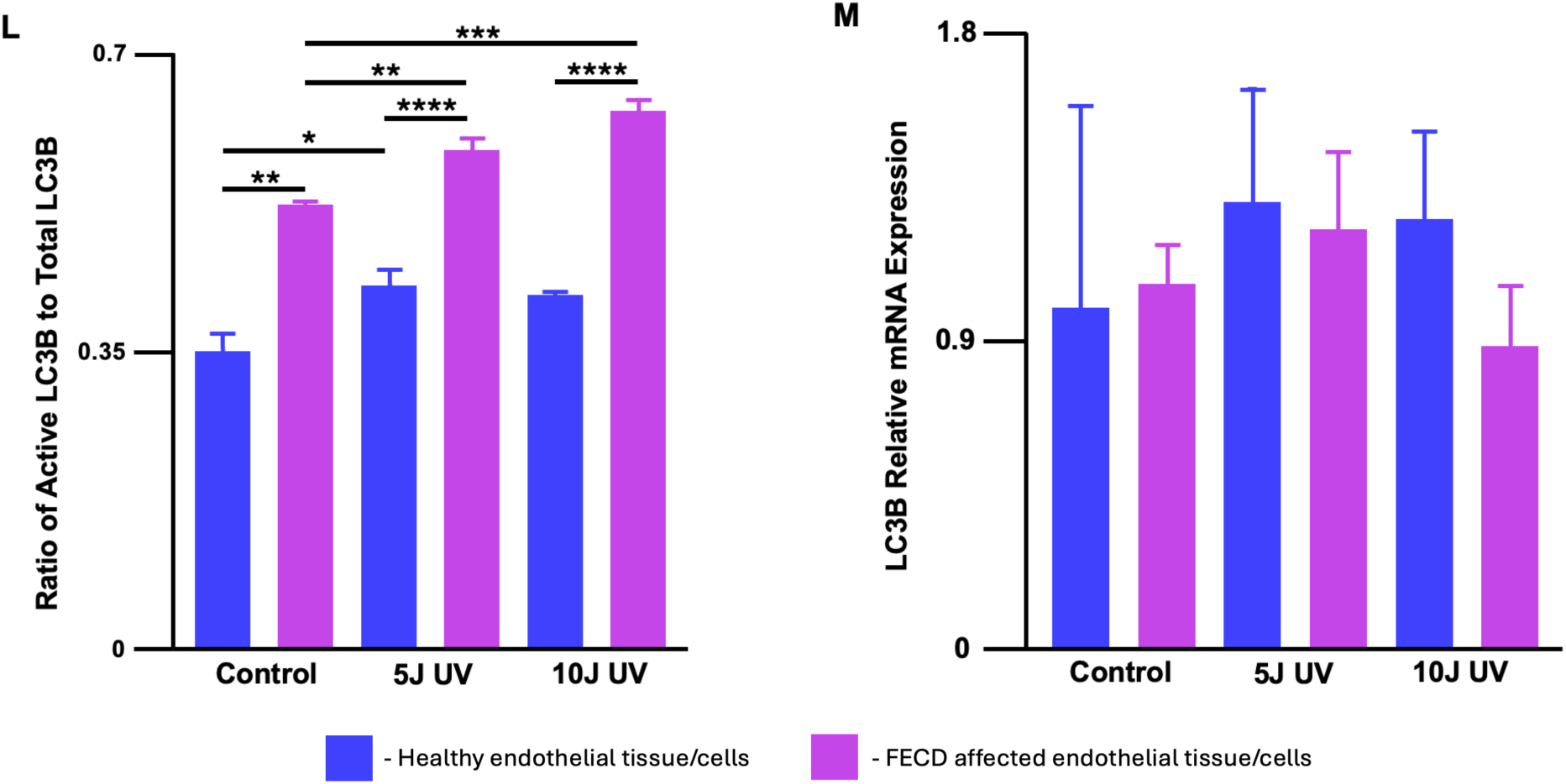
NCOA4 and LC3B expression. (A) NCOA4 protein expression was 2.2-fold higher in surgical FECD cells (N=4) when compared with healthy donor cells (N=8). (B) Immortalized FECD cells (F35T, N=3) showed increased protein expression of NCOA4 when compared with healthy immortalized cells (B4G12, N=3). (C) LC3B activation was increased in immortalized FECD cells (N=4) compared to healthy immortalized control cells (N=4). All data are presented as mean ± SEM; \**P* < 0.05, \*\**P* < 0.01, and \*\*\**P* < 0.001. NCOA4 and LC3B expression. (D) Surgical explant FECD tissue (N=7) showed a 0.43-fold decrease in NCOA4 mRNA levels when compared to healthy donor corneal cells (N=4). (E) There was no significant difference between NCOA4 mRNA levels in immortalized healthy cells (N=3) and immortalized FECD cells (N=3). (F) There was no significant difference between LC3B mRNA levels in immortalized healthy cells (N=3) and immortalized FECD cells (N=3). (G) Immortalized FECD cells (N=3) showed 379-fold production of LC3A mRNA compared with immortalized healthy cels (N=3). All data are presented as mean ± SEM; \**P* < 0.05, \*\**P* < 0.01, and \*\*\**P* < 0.001. NCOA4 and LC3B expression. (H) The amount of LC3B-II (activated) was compared with the total amount of both LC3B-II and LC3B-I (inactivated). The ratio of LC3B activation was greater in FECD vs healthy cells, and this ratio increased further with AMA-induced oxidative damage. (I) Protein levels of ferritinophagy marker NCOA4 increase in both healthy and FECD immortalized cells after undergoing oxidative damage via 100<M AMA exposure. All data are presented as mean ± SEM; \**P* < 0.05, \*\**P* < 0.01, and \*\*\**P* < 0.001. NCOA4 and LC3B expression. (J) 5J UV exposure increased NCOA4 protein expression in both immortalized healthy cells (N=3) and FECD cells (N=3), and 10J UV exposure further increased NCOA4 protein expression in F35T cells. However, these results were statistically insignificant. (K) 5J UV exposure increased NCOA4 RNA expression in both cell lines, and 10J UV exposure further increased RNA expression in B4G12 cells. These results were statistically insignificant. All data are presented as mean ± SEM; \**P* < 0.05, \*\**P* < 0.01, and \*\*\**P* < 0.001. NCOA4 and LC3B expression. (L) Immortalized FECD cells showed increased activation of LC3B autophagic activity at baseline when compared to immortalized healthy cells. UV exposure greatly increased LC3B activation in immortalized FECD cells compared with healthy cells, and greater amounts of UV exposure further increased this disparity. (M) No significant difference was seen in LC3B mRNA expression in healthy or FECD cells undergoing UV exposure. All data are presented as mean ± SEM; \**P* < 0.05, \*\**P* < 0.01, and \*\*\**P* < 0.001.

### FECD cells and healthy cells both express increased ferritinophagy markers with AMA-induced oxidative damage

In order confirm the presence of ferritinophagy in CECs in response to oxidative stress, as well as to determine FECD genotype-specific differences in ferritin sequestration secondary to oxidative stress, AMA was used to induce oxidative damage; NCOA4 protein levels and LC3 autophagy activation were measured as indicators of ferritinophagy. At 500µM AMA treatment, the ratio of LC3B-II to total LC3 increased from 0.45 to 0.81 in FECD immortalized cells (p<0.001) and from 0.31 to 0.56 in immortalized healthy cells (p<0.01), indicating increased autophagy in both cell cultures (**Fig. 1H**). Immortalized FECD cells also showed an 222% protein increase in the ferritinophagy marker NCOA4 when exposed to 50µM AMA to induce oxidative damage (p<0.01), while immortalized healthy cells did not show significantly elevated levels of NCOA4 with AMA exposure (**Fig. 1I**), indicating increased tagging and degradation of ferritin in cells with FECD.

### FECD cells and healthy cells both showed increased ferritinophagy markers with UV exposure

To examine the effect of UVA radiation on ferritinophagy between healthy and FECD affected CECs, immortalized healthy and FECD transformed CECs were exposed to 0J, 5J, and 10J UVA radiation. Protein and mRNA levels of NCOA4 and LC3 were then quantified to assess amounts of ferritinophagy corresponding with differing levels of UV radiation exposure. Immortalized FECD cells had a 7.0% increase in NCOA4 protein production after exposure to 10J UV, and immortalized healthy cells had a 4.5% increase in protein production after 5J UV exposure; however, these results were statistically insignificant (**Fig. 1J**). When exposed to 5J UV radiation, immortalized FECD cells showed increased NCOA4 mRNA (1.48-fold increase), as did immortalized healthy cells (1.43-fold increase); while this data showed positive trends, the results were statistically insignificant (**Fig. 1K)**. No difference was seen in NCOA4 mRNA expression in immortalized FECD cells after 10J UV radiation; after the same 10J UV exposure, NCOA4 mRNA expression in immortalized healthy cells increased 1.53-fold compared to baseline, which was statistically insignificant (**Fig. 1K**). Immortalized FECD cells showed increased LC3B-II activation ratios with 5J UV treatment from 0.52 to 0.59 activation ratio, further increasing to 0.63 LC3B activation after 10J UV exposure (**Fig. 1L**). In healthy immortalized cells, LC3B-II to total LC3 protein ratios increased from 0.35 to 0.43 with 5J UV treatment, with no further increase after 10J UV exposure (**Fig. 1L)**. FECD immortalized cells showed a 1.15-fold increase in LC3B mRNA when treated with 5J UV, dropping to 0.83-fold expression at 10J UV exposure. Healthy immortalized corneal cells showed a 1.31-fold increase in LC3B mRNA when treated with 5J UV, with 1.26-fold expression of LC3B mRNA at 10J UV exposure. These results were statistically insignificant (**Fig 1M)**.

### NCOA4 knockdown increases susceptibility to RSL3-induced ferroptosis in F35T cells and enhances UVA-induced elevation of intracellular ferrous iron (Fe2+)

To investigate the effect of NCOA4 levels on corneal endothelial cell death via ferroptosis, B4G12 and F35T cells were treated with siRNA to decrease NCOA expression. These cells were subsequently treated with RSL3 to induce ferroptosis; control cell samples were treated both with RSL3 as well as with Fer-1 to prevent ferroptosis. B4G12 cells with and without NCOA4 knockdown treatment had no significant differences in cell viability after RSL3 treatment (**Fig 2)**. In F35T cells, cells with NCOA4 knockdown had decreased viability compared with F35T without NCOA4 knockdown (p<0.001). To assess NCOA4’s role in intracellular iron levels in FECD, relative intracellular Fe^2+^ was measured in NCOA4 knockdown and control F35T cells at baseline as well as after 5J UV exposure. No significant difference was seen between Fe^2+^ levels in F35T cells with and without NCOA4 knockdown at baseline; however, after 5J UV treatment, cells with NCOA4 knockdown had significantly higher levels of intracellular Fe^2+^ (p<0.05).

**Figure 2.**
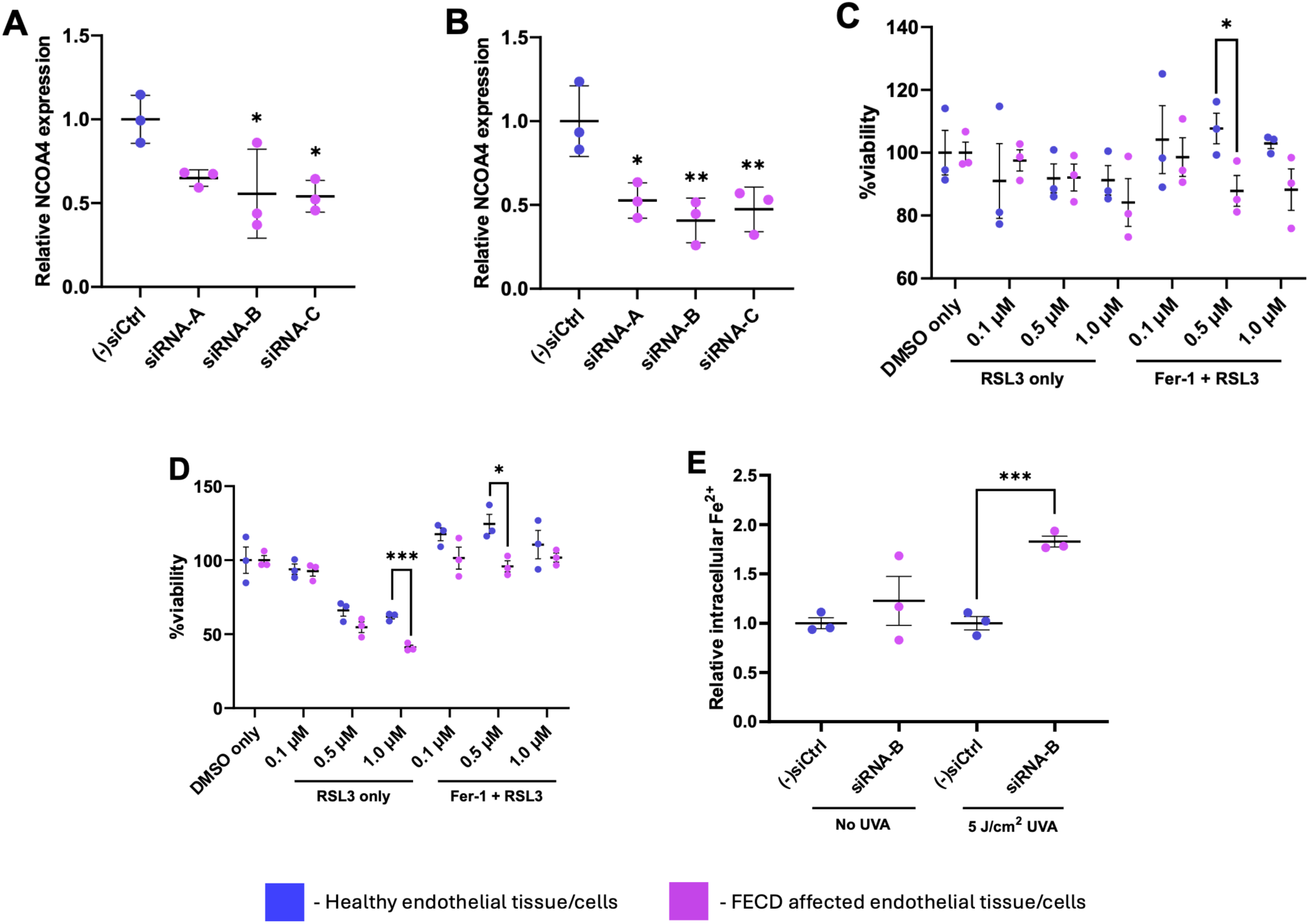
NCOA4 knockdown increases susceptibility to RSL3-induced ferroptosis in HCEC-B4G12 and F35T cells, and enhances UVA-induced elevation of intracellular ferrous ion (Fe^2+^). Relative NCOA4 protein expression levels based on Jess automated western blotting (A) in HCEC-B4G12 and (B) in F35T cells at 72 h after reverse transfection with different anti-NCOA4 siRNAs (siRNA-A, -B, and -C). Post-RSL3 challenge cell viability data (C) in HCEC-B4G12 and (D) in F35T cells with prior NCOA4 knockdown. (E) Relative intracellular ferrous ion (Fe^2+^) in NCOA4-knockdown F35T with or without 5 J/cm^2^ UVA irradiation. All data are presented as mean ± SD; \**P* < 0.05, \*\**P* < 0.01, and \*\*\**P* < 0.001.

### FECD cells showed nuclear localization of ferritin and LC3A

To visually assess healthy donor CECs and surgical explant CECs with FECD for signs of ferritinophagy, cells were tagged with fluorescent markers of NCOA4, ferritin, and LC3. IHC images were obtained from both healthy donor and surgical explant corneas to correspond the observed distribution of ferritinophagy markers with disease burden. Donor cells showed NCOA4, ferritin, and LC3A diffusely located in the cytoplasm of CE cells (**Fig. 3, 4**). Cells from FECD surgical explants showed a pattern of punctate NCOA4 localized in the cytoplasm but showed ferritin and LC3A in the nuclei. This pattern was accentuated for cells closer to the central cornea (in higher diseased states with greater guttae burden) and was lessened in the periphery of diseased corneas (in areas with lower guttae burden).

**Figure 3.**
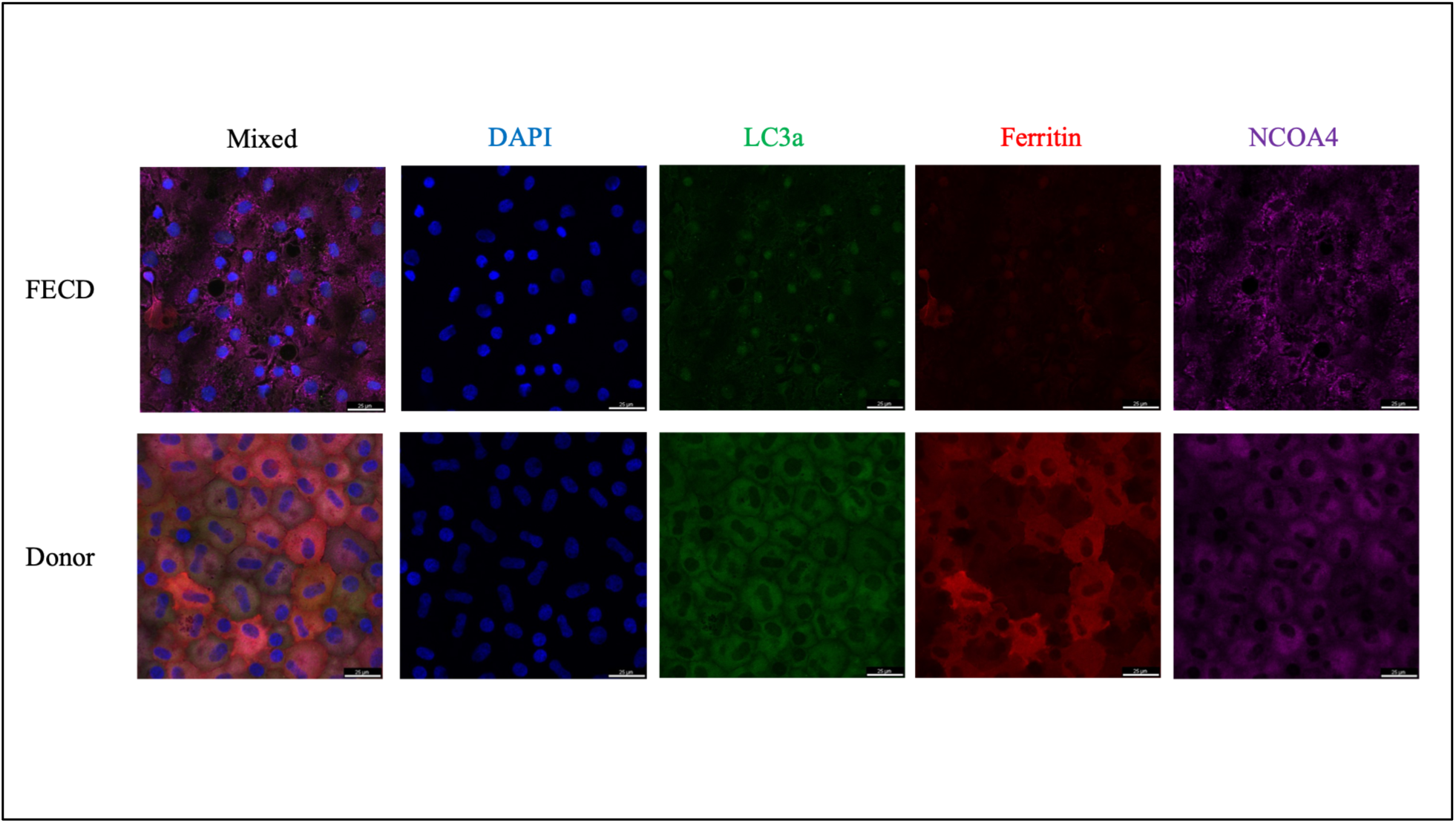
Immunohistochemistry images of FECD cells compared with healthy donor cells. Ferritin (red) decreases dramatically in cells with FECD, indicating the presence of ferritinophagy. NCOA4 and LC3a transition from a diffuse cytosolic appearance in healthy cells to a punctate display in FECD cells, also expected for cells undergoing ferritinophagy. In cells with FECD, ferritin is absent in the cytosol but present in the nuclei, potentially due to a protective mechanism the cell employs to prevent nuclear damage when cellular levels of labile iron increase.

**Figure 4.**
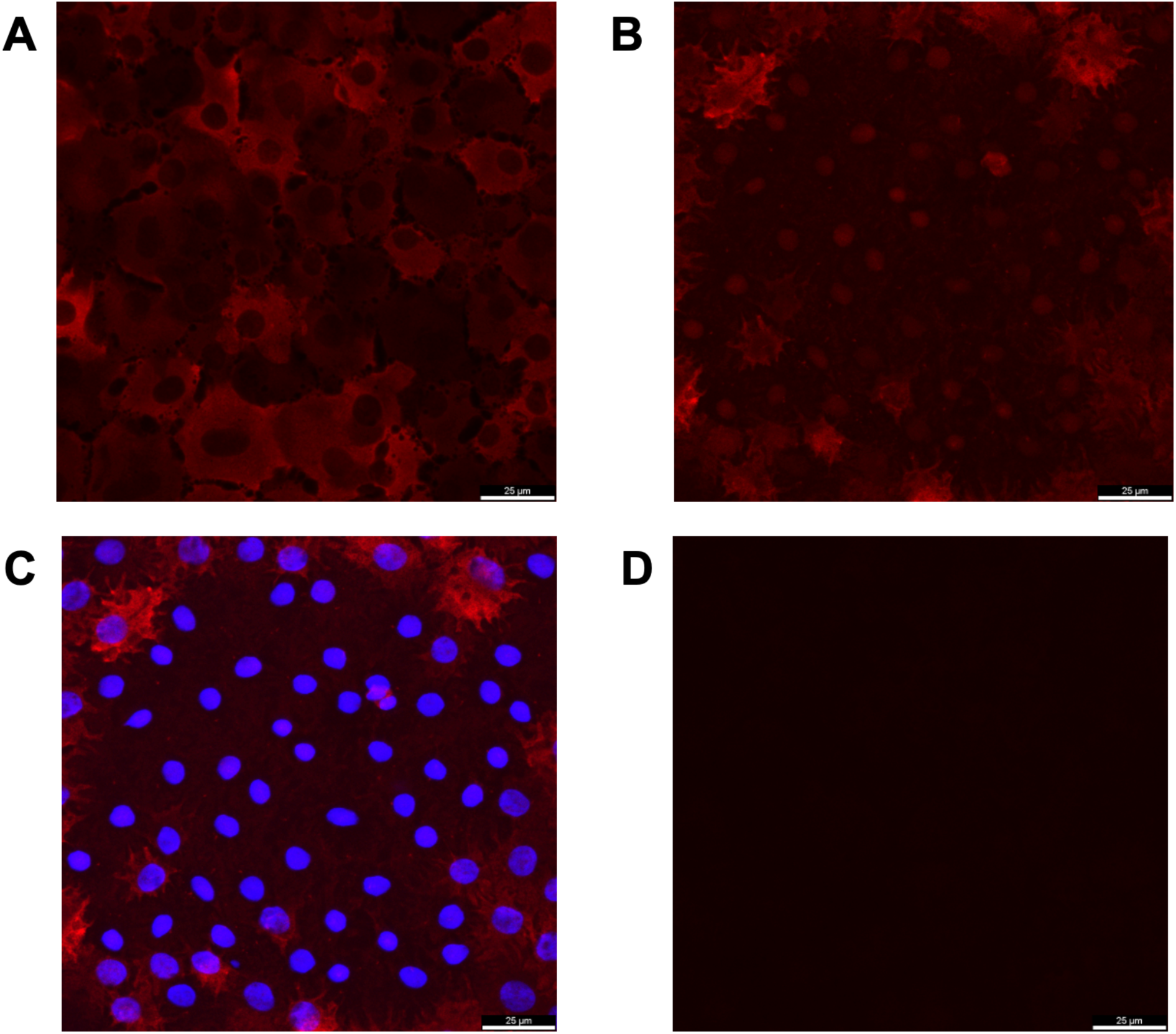
Immunohistochemistry staining of ferritin in healthy donor endothelial cells (A) and endothelial cells collected from corneas with FECD (B). In healthy endothelial cells, ferritin is seen broadly distributed throughout the cytosol; in cells with FECD, ferritin is primarily seen in the nuclei. DAPI staining confirmed ferritin staining in FECD cells correlated with the location of nuclei (C). Negative control confirmed no background staining of ferritin (D).

### FECD corneal endothelial peels demonstrated localization of iron to guttae in iron elemental imaging

To further assess the relationship between iron accumulation and the development of worsening FECD, we utilized LA-ICP-MS to analyze the distribution of iron within corneal peels affected with FECD. When comparing images of endothelial peels under standard microscopy compared to matched images of the same peels after undergoing spectrometry to reveal iron levels, we observed that areas with guttae formation (indicative of worsened FECD) had higher iron intensity levels (**Fig. 5**). These results were observed in three separate corneal endothelial peels, all of which displayed localization of higher iron levels in areas of guttae formation.

**Figure 5.**
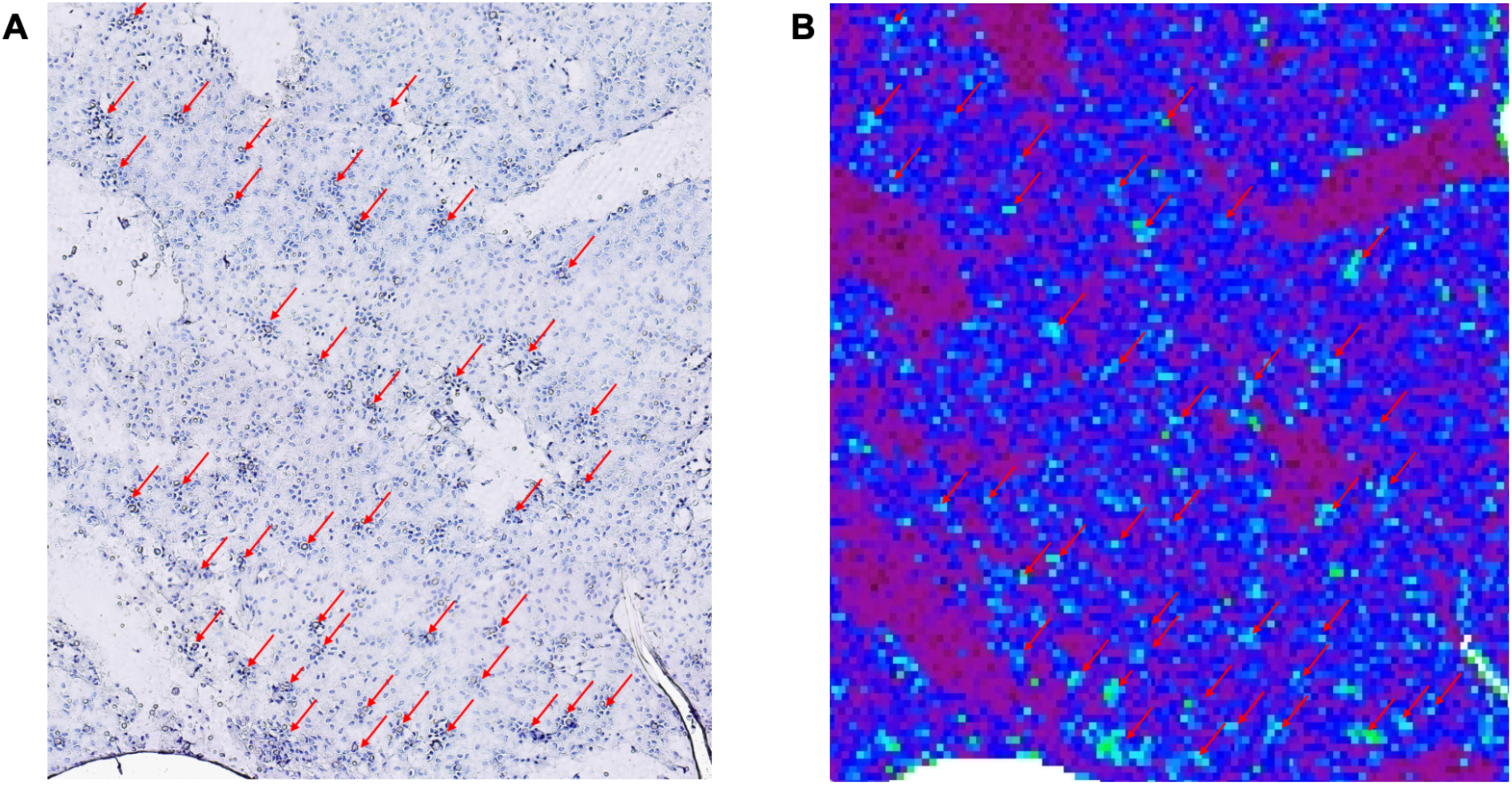
Iron Spec images showing a correlation with guttae formation on endothelial imaging (A) with increased iron levels under spectrometry (B). Increased iron levels are delineated by green or light-blue pixels. Red arrows highlight guttae.

## DISCUSSION

In cells, ferritin executes the important function of iron storage, as each ferritin protein complex can contain up to 4,500 iron atoms [15]. Ferritin is composed of 24 subunits of light chains (FTL) and heavy chains (FTH). Of these subunits, heavy chain 1 (FTH1) serves an important role in the process of ferritinophagy, the process by which ferritin is degraded to release iron intracellularly [11]. Ferritinophagy begins with the formation of the autophagosome, which is mediated by the ATG family of proteins [24]. ATG4 cleaves protein LC3 to form LC3-I, and when autophagy is initiated, LC3-I becomes bound to phosphatidylethanolamine (PE) on the phagophore’s membrane. This newly attached protein is termed LC3-II, and because it remains anchored to the autophagosome, it serves as a common molecular marker to observe autophagy in cells. During autophagy, fluorescently tagged LC3-II appears aggregated in a punctuated pattern, reflecting its conjugation to PE on the autophagosome membrane. The LC3 family has several variants; LC3A specifically has been found to localize to the autophagosome membrane and is expressed in almost all tissues, and was thus used as the autophagy marker for some of our experiments [28], although LC3B was also used for some of our protein quantification experiments based on its high expression observed in western blots in CEC samples. By detecting increased levels of LC3-II compared with total LC3 in CECs with FECD, this study confirms the presence of increased autophagy FECD. Our previous work revealed increased intracellular Fe^2+^ accumulation leading to ferroptosis in CECs with FECD [5], so we examined the additional biomarker NCOA4 to directly connect the increased autophagy and ferroptosis observed with ferritinophagy.

Ferritinophagy occurs naturally in the cell when iron levels decrease, allowing the cell to degrade ferritin to release more iron. In the process of ferritinophagy, NCOA4 – a transcriptional coactivator for several nuclear receptors including androgen receptor and estrogen receptor alpha that is found in both the nucleus and the cytosol [29, 30] – binds to the FTH1 region of ferritin, targeting it for degradation and the release of iron into the cytosol [11, 31]. On the other hand, ferritinophagy becomes inactivated when cellular iron levels rise, whereby NCOA4 binds to HERC2 and is subsequently degraded to prevent further ferritin degradation and release of iron into the cytosol. In ferritinophagy, once NCOA4 has become bound to FTH1, ferritin is delivered to the autophagosome and ultimately causes ferritin degradation. Because of the central role of NCOA4 in ferritinophagy, it serves as another useful biomarker for ferritin degradation; the interplay between NCOA4 and ferritin can be observed to insinuate ongoing or recent ferritinophagic activity. When LC3, NCOA4, and ferritin colocalize in puncta, the process of ferritinophagy is likely actively occurring [32]. On the other hand, high levels of punctate cytosolic NCOA4 with low levels of cytosolic ferritin indicate a recently post-ferritinophagic state, as the ferritin has already been degraded, but the residual autophagosomes have yet to be cleared. Importantly, ferritinophagy is tightly connected with cell death via ferroptosis. Decreasing levels of NCOA4 has been shown to repress ferritin degradation, thus lowering the occurrence of ferroptosis [19]. By tracking the occurrence and flux of ferritinophagy in several models of FECD including patient derived surgical samples and human FECD cultured cells with the well-characterized and common *TCF4* trinucleotide repeat expansion, this investigation can provide powerful insights into the mechanism that results in CEC death and better inform our understanding of disease progression.

In our experiments, we were able to demonstrate increased ferritinophagy in CECs with FECD by showing both increased levels of NCOA4 expression and increased LC3-II activation relative to total LC3, indicating active tagging and autophagy of ferritin. The upregulation and increased expression of key ferritinophagy proteins in FECD opens an additional explanation for the oxidative damage and cellular death seen in CECs affected by this disease. In ferroptosis, the production and accumulation of lipid peroxides results in a non-apoptotic form of cell death [33]. When there is free iron available intracellularly, these lipid peroxides can be created by the interaction of polyunsaturated fatty acids (PUFAs) with hydroxyl radicals formed via Fenton reactions [34]. Additionally, iron-dependent lipoxygenases can contribute to the formation of lipid peroxides [33, 35]. Both of these methods rely on an abundance of intracellular iron, which, as discussed above, can be released from ferritin through ferritinophagy. The increase in ferritinophagy proteins and markers observed in CECs with FECD provides solid evidence that ferritinophagy mediates intracellular iron accumulation and resultant ferroptosis in FECD [36].

Our investigation also provides evidence that oxidative stress in CECs, particularly FECD CECs exposed to UVA light, results in increased ferritinophagic activity. Previous studies have established that UV exposure increases DNA damage in FECD, contributing to the disease’s degradative process [9]. CECs affected with FECD are particularly susceptible to sources of oxidative stress including UVA, which impacts all layers of the cornea by inducing the production of ROS intracellularly [37, 38]. Most of this UV light exposure occurs in the central cornea at the apex of the eye, with attenuation of exposure in the periphery in part from the effects of collagen scattering [39]. This asymmetry mirrors the central to peripheral spread of CEC loss and guttae formation seen in corneas with FECD, supporting the evidence that UVA light exposure drives FECD progression [9]. Our experiment complements this knowledge by demonstrating that UVA exposure also increases the rate of ferritinophagy in cells with FECD, furthering the oxidative damage and cellular death by ferroptosis that results from cytosolic iron accumulation as our group demonstrated previously [5]. CECs with and without FECD both displayed higher degrees of LC3B activation after undergoing exposure to UVA, indicating increased autophagic activity. While we were unable to obtain significant results regarding increasing levels of NCOA4 protein levels in CECs undergoing UVA exposure, our initially data trends show an increase in NCOA4 protein levels in CECs with FECD that have been exposed to UVA. We believe that with further replications and a higher number of treatment samples, these preliminary trends will show statistical significance. To further assess the connection between oxidative stress and ferritinophagy, our study utilized AMA as an alternative source of inducing oxidative damage in cell cultures and tissues. AMA contains a formamidosalicylic acid functional group that allows the molecule to inhibit multiple components of the electron transport chain, including cytochrome b, cytochrome c, succinate oxidase, and NADH oxidase [23, 40]. This causes electrons to leak out of the ETC system, eventually resulting in ROS formation. This ETC inhibition also destroys the proton gradient across the inner mitochondrial membrane, decreasing the mitochondrial membrane potential [41]. Our results show that AMA induced oxidative damage mimic the results seen in UV treated cells; cells treated with AMA show increases in NCOA4 expression and LC3 activation, indicative of increased ferritinophagy. This indicates that UV exposure likely causes ferritinophagy secondary to the induction of oxidative damage, rather than via a UV specific mechanism. Taken altogether, AMA and UVA oxidative stress exposure results reinforces the important role of ferritinophagy in FECD disease and provide important future areas to develop deeper understanding of FECD disease progression.

To further substantiate the connection between iron accumulation and FECD progression, we analyzed corneal peels under microscopy as well as with LA-ICP-MS to correlate FECD progression (as evidenced by loss of endothelial architecture and guttae formation) with the presence of higher levels of iron accumulation. When we compared spectrometry images with endothelial peel images under microscopy, we observed that higher levels of iron were clearly observed in the same spatial arrangement as guttae (**Fig. 5**). Once again, this would be expected with our proposed mechanism of ferritinophagy in FECD. As FECD progresses, cells with worsened disease states will cause disruptions to the endothelial layer architecture and clump together; if ferritinophagy was uncontrollably occurring in these cells, we would expect to observe higher levels of iron in areas of higher FECD burden. Given that we were able to use the same corneal peels under general microscopy as well as under spectrometry, we were able to correlate areas of higher FECD activity with higher levels of iron accumulation, further substantiating the role of ferritinophagy and ferroptosis in FECD.

Some unexpected results were observed in cells with NCOA4 knockdown that had been treated with RSL3 to induce ferroptosis. Because of NCOA4’s role in tagging ferritin for degradation, we expected that NCOA4 knockdown would increase cell viability in cells with induced ferroptosis; in our experiments, NCOA4 knockdown F35T cells had decreased rather than increased viability. Gryzik et al. found similar results in their research, which investigated the role of NCOA4 in cells with induced ferroptosis [42]. Notably, Gryzik et al. induced ferroptosis through two different methods: erastin exposure, and RSL3 exposure. They found that while NCOA4 knockdown was protective in cells undergoing ferroptosis via erastin exposure, cells with NCOA4 knockdown experienced decreased survival rates compared to control cells when treated with RSL3 to induce ferroptosis. This may indicate that RSL3-induced ferroptosis is less dependent on ferritin degradation when compared to erastin-induced ferroptosis. Additionally, while NCOA4 has been implicated in ferritin tagging for degradation, its full mechanisms are not fully understood. It is possible that NCOA4 plays a separate, protective role in ferroptosis unrelated to ferritin tagging and degradation, which would explain the negative impact of NCOA4 knockdown on viability during RSL3-induced ferroptosis. Our results showed that in FECD CECs undergoing UV exposure, NCOA4 knockdown increased the amount of intracellular Fe^2+^. Once again, this result seems to indicate that the role of NCOA4 is more multifaceted than previously assumed; while it certainly contributes to ferritin degradation and iron accumulation via ferritinophagy, it may also function to decrease iron burden, particularly in cells undergoing oxidative stress.

While the presence of increased ferritinophagy activity in FECD has been demonstrated by increased levels of NCOA4 and LC3 in CECs, the images we obtained through immunohistochemistry were somewhat unexpected. The expected appearance of active ferritinophagy is the colocalization of ferritin, NCOA4, and LC3A into puncta found in the cytosol of the cell. As expected, in healthy donor CE cells, NCOA4, LC3A, and ferritin were observed throughout the cytosol. Cells from surgical explant tissues affected by FECD displayed a pattern indicative of post-ferritinophagy: NCOA4 was seen throughout the cytosol in a punctate distribution, and ferritin was absent from the cytosol. However, we noted an unanticipated finding on IHC: both ferritin and LC3A localized inside the nuclei of affected cells. LC3 can enter the nuclei of cells via passive diffusion, and when cells are primed for autophagy (such as during starvation states), LC3 is deacetylated to allow for facilitated transport out of the nucleus into the cytosol [43]. It is possible that transport of LC3 out of the nucleus is inhibited during FECD, either by the disease itself or by some protective mechanism of the cell to decrease ferritinophagy. Ferritin has been observed in the nuclei of corneal epithelial cells in chicken embryos [44]. It is assumed that ferritin plays a protective role in these cells by sequestering any iron that has diffused into the nuclei, thereby preventing Fenton reactions from occurring close to the DNA. This is particularly important in cells with high UV exposure, as seen in the corneal epithelium. The transport of ferritin into the nuclei of these cells is accomplished by a protein called ferritoid [45]. We hypothesize that in FECD, the excess of iron released during ferritinophagy diffuses into the nucleus, triggering an increase of ferritoid production. This in turn shuttles ferritin into the nucleus in an attempt to sequester the excess iron away from DNA (**Fig. 6**). By separating ferritin from NCOA4, ferritinophagy would necessarily decrease, further protecting the cell. Additionally, we suspect that CECs attempt cell rescue during excess ferritinophagy by translocating LC3 into the nuclei of the cells; this would provide one more layer of protection by keeping a critical protein for autophagy away from the cytosol. While the full mechanism of ferroptosis has not been completely elucidated, this insight has the potential to open new avenues of FECD disease treatment, as targeting ferroptosis and ferritinophagy may prevent the degradative progression of FECD.

**Figure 6.**
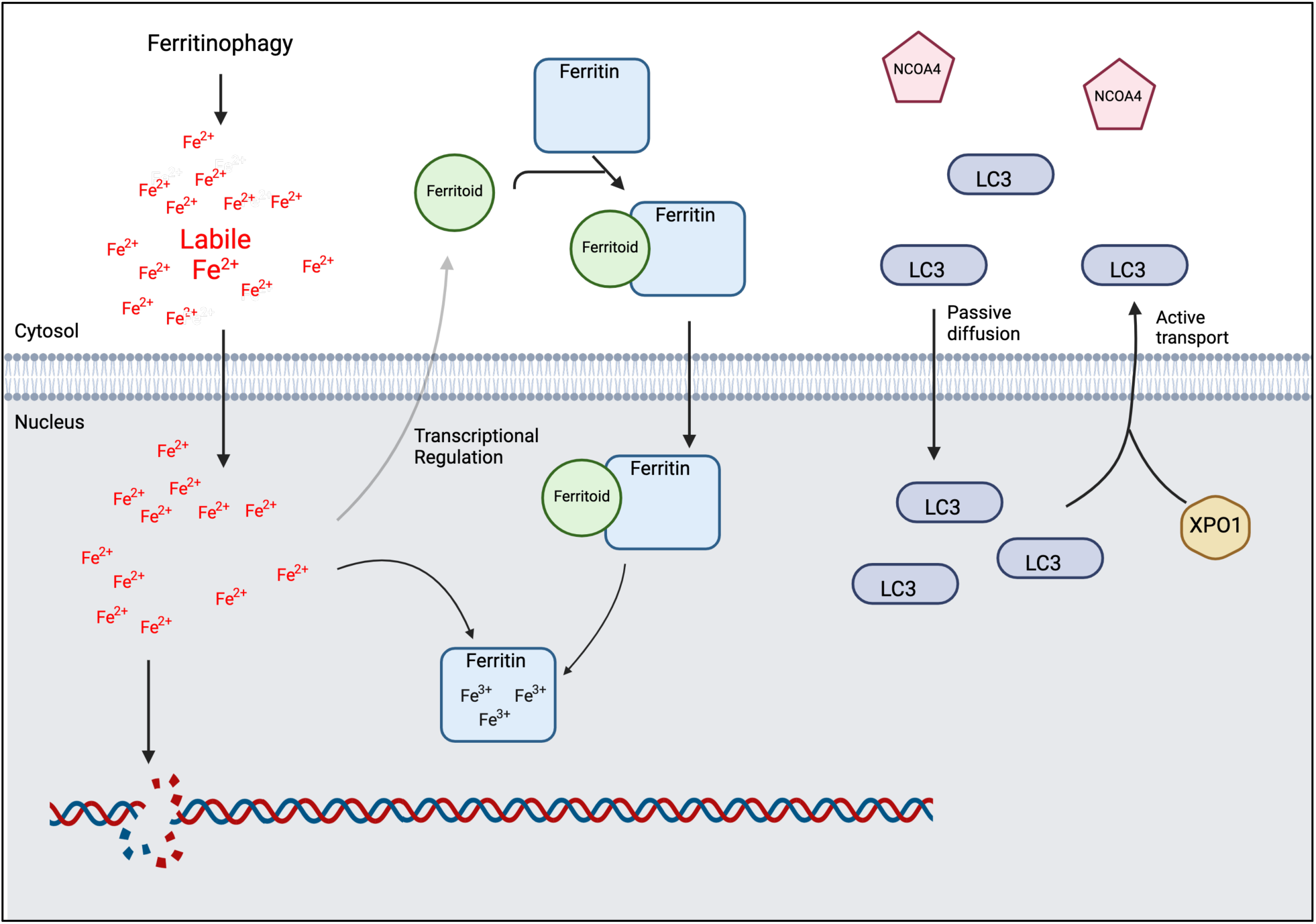
Proposed mechanism for nuclear ferritin. As seen in Figure 3, ferritin in cells with FECD is exclusively found in the nuclei. This potentially results from a protective mechanism by which a transport protein shuttles ferritin from the cytosol to the nucleus in response to high labile iron levels. This protects the cell by isolating nuclear iron into ferritin, thereby stopping Fenton reactions and subsequent oxidative damage from damaging nuclear DNA.

There are additional studies that we propose to further elucidate the important role that ferritinophagy plays in FECD pathogenesis. We propose investigating the possibility that ferritoid functions as a nuclear transporter of ferritin in CECs with FECD. If present, this would confirm a self-defense mechanism exists in cells against iron-induced damage in FECD. We also propose continued investigation of LC3A and LC3B in health and diseased CECs to rule out disparate expression of either protein subtype in ferritinophagy in FECD. Lastly, we would like to replicate the results of our UV exposure experiments with surgical and donor tissues. Our results from UV studies have been obtained primarily from immortalized cell lines, and it is important to conduct UV experiments in surgical tissues with donor corneal tissue controls, which was outside the scope of this current investigation.

In conclusion, this experiment furthers the current understanding of FECD by specifically implicating the process of ferritinophagy with the previously examined Fe^2+^ accumulation noted in CECs with FECD. Moreover, our work connects the rise of ferritinophagy in FECD with UVA radiation exposure in CECs, providing a plausible mechanism for the negative effects of UV exposure on FECD progression. Lastly, IHC imaging shows evidence of post-ferritinophagic changes in CECS with FECD, revealing cellular rescue attempts in the translocation of both ferritin and LC3 into the nuclei of affected CECs. Taken together, these results demonstrate the role of ferritinophagy in FECD pathogenesis and provide potential biomarkers to target for further therapeutic developments.

## Scope

This study investigates the source of aberrant iron in Fuchs endothelial corneal dystrophy and uses cell models and human tissue to study the effects of genetic background and environmental factor UVA on disease pathology.

## Acknowledgments

This research was made possible by generous support from Mr. and Mrs. Robert and Joell Brightfelt, Mr. and Mrs. Lloyd and Betty Schermer, Ms. Mary Dawson, the M.D. Wagoner & M.A. Greiner Cornea Excellence Fund, the Cornea Transplant Research Fund, the Beulah and Florence Usher Chair in Cornea/External Disease and Refractive Surgery, the Seidler Educational Support Fund in Cornea Research, the National Eye Institute, and our patients, cornea donors, and donor families. The authors would also like to thank Apurva Dusane, M.Pharm., for technical assistance.

